# Happy Cow or Thinking Pig? WUR Wolf – Facial Coding Platform for Measuring Emotions in Farm Animals

**DOI:** 10.1101/2021.04.09.439122

**Authors:** Suresh Neethirajan

## Abstract

Emotions play an indicative and informative role in the investigation of farm animal behaviors. Systems that respond and can measure emotions provide a natural user interface in enabling the digitalization of animal welfare platforms. The faces of farm animals can be one of the richest channels for expressing emotions. We present WUR Wolf (Wageningen University & Research: Wolf Mascot)—a real-time facial expression recognition platform that can automatically code the emotions of farm animals. Using Python-based algorithms, we detect and track the facial features of cows and pigs, analyze the appearance, ear postures, and eye white regions, and correlate with the mental/emotional states of the farm animals. The system is trained on dataset of facial features of images of the farm animals collected in over 6 farms and has been optimized to operate with an average accuracy of 85%. From these, we infer the emotional states of animals in real time. The software detects 13 facial actions and 9 emotional states, including whether the animal is aggressive, calm, or neutral. A real-time emotion recognition system based on YoloV3, and Faster YoloV4-based facial detection platform and an ensemble Convolutional Neural Networks (RCNN) is presented. Detecting expressions of farm animals simultaneously in real time makes many new interfaces for automated decision-making tools possible for livestock farmers. Emotions sensing offers a vast amount of potential for improving animal welfare and animal-human interactions.

## 1. Introduction

Digital technologies, in particular, precision livestock farming, and artificial intelligence have the potential to shape the transformation in animal welfare [1]. To ensure access to sustainable and high-quality health attention and welfare in animal husbandry management, innovative tools are needed. Unlocking the full potential of automated measurement of mental and emotional states of farm animals through digitalization such as facial coding systems would help to blur the lines between biological, physical, and digital technologies.

Animal caretakers, handlers, and farmworkers typically rely on hands-on observations and measurements while investigating methods to monitor animal welfare. To avoid the increased handling of animals in the process of taking functional or physiological data, and to reduce the subjectivity associated with manual assessments, automated animal behavior and physiology measurement systems can complement the current traditional welfare assessment tools and processes in enhancing the detection of animals in distress or pain in the barn [2]. Automated and continuous monitoring of animal welfare through digital alerting is rapidly becoming a reality [3].

In the human context, facial recognition platforms have long been in use for various applications, such as password systems on smartphones, identification at international border checkpoints, identification of criminals [4, diagnosis of Turner syndrome [5]; detection of genetic disorder phenotypes [6]; as a potential diagnostic tool for Parkinson disease [7]; measuring tourist satisfaction through emotional expressions [8]; and quantification of customer interest during shopping [9].

### Emotions

Emotions are believed to be a social and survival mechanism that is present in many species. In humans, emotions are understood as deep and complex psychological experiences that influence physical reactions. There is an entire sector of science devoted to understanding the sophisticated inner workings of the human brain, yet many questions related to human emotions remain unanswered. Even less scientific research is focused on understanding the emotional capacity of non-human primates and other animals. The ability to interpret the emotional states of an animal is considerably more difficult than understanding the emotional state of a human [10]. The human face is capable of a wide array of expressions that communicate emotion and social intent to other humans. These expressions are so clear that even some non-human species, like dogs, can identify human emotion through facial expression [11]. Each species has its own unique physiological composition resulting in special forms of expression. Despite human intellectual capacity, emotional understanding of other species through facial observation has proven difficult.

Early studies [12] noted the influence of human bias on interpretations and accidental interference with the natural responses of animals. It is not uncommon for humans to anthropomorphize the expressions of animals. The baring of teeth is an example. Humans commonly consider such an expression to be “smiling” and interpret it as a sign of positive emotions. In other species, such as non-human primates, the baring of teeth is more commonly an expression of a negative emotion associated with aggression [13].

For these reasons and many others, the involvement of technology is critical in maintaining accurate and unbiased assessments of animal emotions and individual animal identification. In recent years, the number of studies concerning technological intervention in the field of animal behavior has increased [10]. The ability of customized software to improve research, animal welfare, the production of food animals, legal identification, and medical practices is astounding.

### Understanding Animal Emotions

The human comprehension of animal emotions may seem trivial; however, it is a mutually beneficial skill. The ability of animals to express complex emotions, such as love and joy, is still being debated within the field of behavioral science. Other emotions, such as fear, stress, and pleasure are studied more commonly. These basic emotions have an impact on how animals feel about their environment and interact with it. It also impacts an animal’s interactions with its counter specifics.

Non-domesticated species of animals are commonly observed in the wild and maintained in captivity to understand and conserve their species. Changes in the natural environment, because of human actions, can be stressful for individuals within a species. Captive non-domesticated animals also experience stress created through artificial environments and artificial mate selection. If even one animal experiences and displays signs of stress or aggression, its companions are likely to understand and attempt to respond to their emotional state [14]. These responses can result in stress, conflict, and the uneven distribution of resources [15]. The understanding of emotional expression in captive animals can help caretakers determine the most beneficial forms of care and companion matching for each individual, resulting in a better quality of life for the animals in question.

Companion animals are another category of individuals who can benefit from a deeper understanding of animal emotion. Just like humans, individual animals experience different thresholds for coping with pain and discomfort. Since many companion animals must undergo voluntary medical procedures for the well-being of their health and their species, it is important to understand their physical responses. Animals cannot tell humans how much pain they are in, so it is up to their caretakers to interpret the pain level an animal is experiencing and treat it appropriately [16]. This task is most accurately completed when the emotions of an animal are clearly and quickly detectable.

The understanding of expressions related to stress and pain is impactful in animal agriculture. Animals used for food production often produce higher quality products when they do not experience unpleasant emotions [17]. The detection of individual animals experiencing stress allows for the early identification of medical complications as well. A study on sows in parturition showed a uniform pattern of facially expressed discomfort during the birthing cycle [18]. In such a case, facial identification of emotional distress could be used to detect abnormally high levels of discomfort and alert human caretakers to the possibility of dystocia.

### Facial Recognition Software

Facial recognition software has been used on human subjects for years. It has even contributed to the special effect capabilities in films and is used as a password system for locked personal devices. It is a non-invasive method that tracks specific points on an individual’s face using photos and videos. These points need not be placed directly on the subject’s face; instead, computer software can be customized and trained to identify the location of each point. Once this software identifies an individual’s characteristics, it can be modified to detect changes in facial positioning and associate those changes with emotional states.

The same method can be used to identify individuals and emotional states when it comes to animal subjects. With a bit of software reconstruction, scientists have been able to create reliable systems for the assessment of animal emotions through technological means. These systems have been specified to identify multiple species including, cows, cats, sheep, large carnivores, and many species of non-human primate. In studies focused on identifying individual members of the same species within a group, the accuracy of specialized facial recognition software was found to be between 94% and 98.7%. Some of these studies even displayed the ability of software to identify and categorize new individuals within a group and the ability to identify individuals at night [19, 20, 21]. Other studies focused more on the emotional expressions that could be identified through facial recognition software and some of the studies showed an accuracy of around 80% when compared to the findings of professionals in the field of animal emotion identification [22].

### The Grimace Scale

The facial recognition software used is based on a series of points in relation to phenotypic features of the species in question, but it uses an older theory to attach the location of those points to emotional states.

The grimace scale is a template created to depict the physical reactions associated with varying levels of discomfort. These scales are created in relation to a specific species and are defined by a numerical scale [23]. In the case of pigs, sheep, and cattle, grimace scales normally focus on tension in the neck, shape of the eye, tension in the brow, nose bunching, and positioning of the ears [24, 18]. These visual cues can be combined with vocal cues to further depict the level of discomfort an animal is experiencing. In species like mice, other expressive physical features must be accounted for, such as whisker movement [25]. For less social species, like cats, the changes in facial expression in response to pain are more minute but still identifiable with the use of a grimace scale [16].

These scales have been proven as an accurate way to assess pain with minimal human bias [23]. They are created through the professional observation of species during controlled procedures that are known to trigger pain receptors. Once created, grimace scales can be converted to specific measurements that are detectable through facial recognition software with the assistance of the Viola-Jones algorithm. This algorithm breaks down the facial structure of animal subjects into multiple sections to refine, crop, and identify major facial features [22]. These features make the technological interpretation of animal emotions feasible across a variety of species and in a variety of settings.

### Best Way to Manage Animal Emotion Recognition

Studies are most accurate when the spectrum of discomfort, including everything from acute low-grade pain to severe chronic pain, is fully identified. Events of low-grade discomfort are significant; however, they may not be identifiable through the production of the stress hormone cortisol [26]. In such situations, discomfort may only be discernable through the facial expressions of an animal detectable by facial recognition software.

On large-scale farms, it is important to keep the animals comfortable and relaxed, but it would be impractical and expensive to test the chemical levels of stress present in every animal. The identification of emotional states through facial recognition software provides a more efficient and cost-effective answer. It also provides an opportunity for the identification of very similar individuals in a way that cannot be illegally altered, unlike ear tags, which are sometimes changed for false insurance claims [21].

The use of facial recognition software also reduces the need for human interaction with animal subjects. For non-domesticated animals, the presence of human observers can be a stressful experience and alter their natural behavior. Facial recognition software allows researchers to review high-quality video and photo evidence of the subject’s emotional expressions without any disturbance. Researchers can even record the identification and actions of multiple individuals within a group of animals at the same time with the help of software like LemurFaceID [20].

Room for human error in the form of bias is reduced with the help of facial recognition software. Since humans’ experience emotions and have the ability to empathize with other emotional beings, human observers run the risk of interpreting animals’ emotional expressions improperly. In a study concerning the pain expressions of sows during parturition, it was noted that all female observers rated the sows’ pain significantly higher than the male observers [18]. That is not to say that one gender is more capable of emotional recognition than the other; rather, this situation highlights the opportunity for human error and bias when assessing animals’ emotions. With well-calculated software, these discrepancies will cease to exist, and researchers can focus more of their time on finding significant points and connections within recorded data, rather than spending their time recording the data.

## 2. Materials and Methods

### Dataset

Images (Figure 1) and videos of cows and pigs were collected from multiple locations: 3 farms in Canada, 2 farms in the USA, and 1 farm in India. The dataset consisted of 3780 images from a total of 235 pigs and 210 dairy cows. These images were grouped and categorized into multiple subfolders, based on 3 emotions of cows and 6 emotions of pigs. The farm animal’s facial expression started from positive to neutral to negative states and returns to neutral state during the data collection process. No elicitation or inducement of affective states on the farm animals were conducted during the data collection.

**Figure 1.**
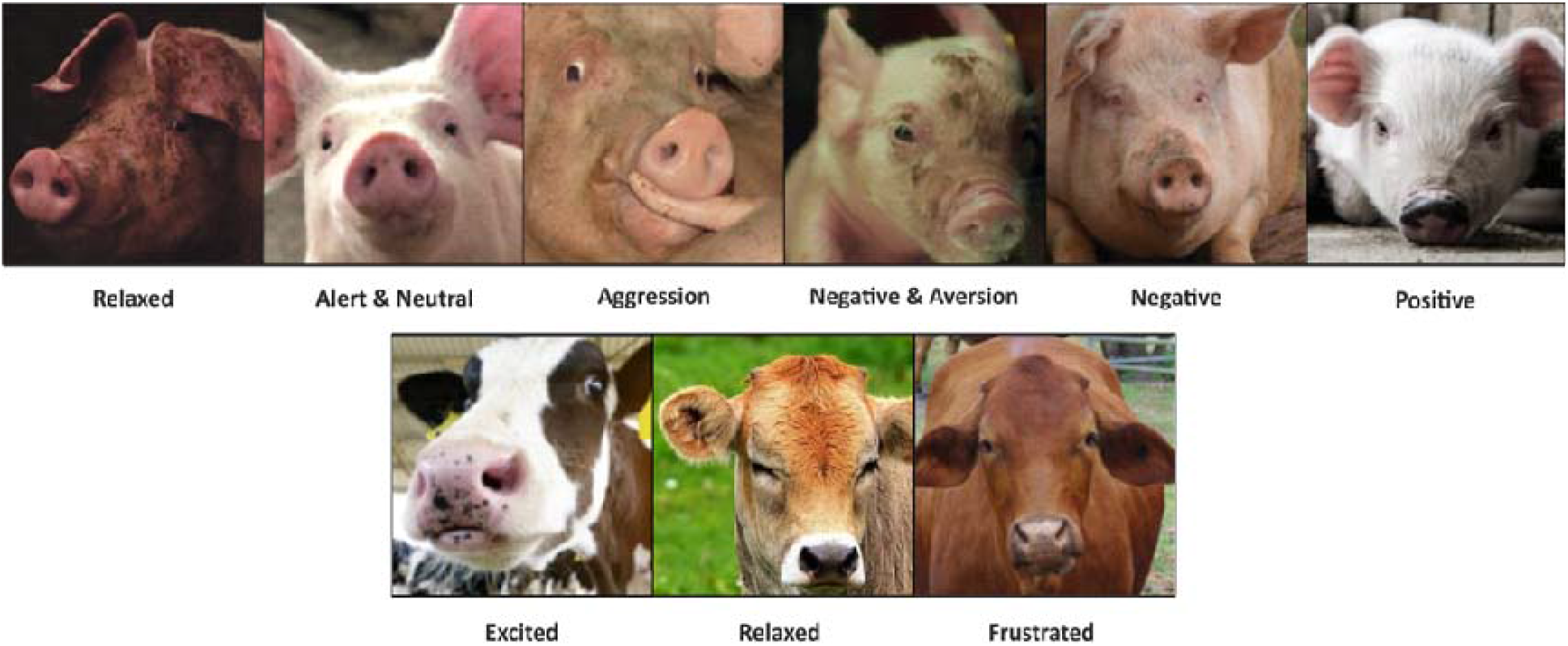
Sample of images from the data set. Facial features of pigs and cows expressing varying emotions.

#### Features and Data Processing

The collected and grouped images dataset were divided into 9 classes based on the correlation between the facial features such as ear posture and eye whites of cows and pigs and the sensing parameters as compiled in Table 1. The videos and images were preprocessed initially using a 3-stage method (1) Detection of faces, (2) Alignment of faces, (3) Normalization of input. Regular smartphone (Samsung Galaxy S10) was used for capturing images and videos from different angles and directions when the animals were in the barn or pen. The collected data were labelled based on the time stamp and the RFID tags and markers. Faces were not manually extracted but by the MIT LabelImg code [27]. Annotations for labeling different models’ bounding boxes were done in the standard format for each: PASCAL format for Faster-RCNN and YOLO format for both YOLOv3 and YOLOv4.

**Table 1.** Sensing parameters that were used for each of the 9 classes related to recognizing emotions of cows and pigs [10].

| Species Type | Indicators Inferring Emotions | Emotions/Affective States |
| --- | --- | --- |
| Cow | Upright ear posture longer | Excited state |
| Cow | Forward facing ear posture | Frustration (negative emotion) |
| Cow | Half-closed eyes and ears backwards or hung-down | Relaxed state |
| Cow | Eye white clearly visible and ears directed forward | Excited state |
| Cow | Visible eye white | Stress (negative emotion) |
| Pigs | High frequency ear movement | Stress (negative emotion) |
| Pigs | Ears forward | Alert (neutral emotion) |
| Pigs | Ears backward | Negative emotion |
| Pigs | Hanging ears flipping in the direction of eyes | Neutral emotion |
| Pigs | Standing upright ears | Normal (neutral state) |
| Pigs | Ears forward oriented | Aggression (negative emotion) |
| Pigs | Ears backward and less open eyes | Retreat from aggression or transition to neutral state |

#### Hardware

The training and the testing of the 3 models based on YoloV3, YoloV4 and Faster RCNN were performed on NVidia GeForce GTX 1080 Ti graphics processing unit (GPU) running on CUDA 9.0 (compute unified device architecture) and CUDANN 7.6.1 (CUDA deep neural network library), equipped with 3584 CUDA cores and 11 GB memory.

### YOLOv3

You Only Look Once (YOLO) is one of the fastest Object Detection Systems with a 30 FPS image processing capability and a 57.9% mAP (mean Average Precision) score [28]. YOLO is based on a single Convolutional Neural Network (CNN), i.e. one-step detection and classification. The CNN divides an image into blocks and then it predicts the bounding boxes and probabilities for each block. It was built on a custom Darknet architecture: darknet-19, a 19-layer network supplemented with 11 object detection layers. This architecture, however, struggled with small object detections. YOLOv3 uses a variant of Darknet, a 53-layer Imagenet - trained network combined with 53 more layers for detection and 61.5M parameters. Detection is done at 3 receptive fields: 85 × 85, 181 × 181, 365 × 365, addressing the small object detection issue. The loss function doesn’t utilize exhaustive candidate regions but generates the bounding box coordinates and confidence using regression. This gives faster and more accurate detection. It consists of 4 parts, each given equal weightage: regression loss, confidence loss, classification loss, and loss for the absence of any object. When applied to face detection, multiple pyramid pooling layers capture high-level semantic features, and the loss function is altered. Regression loss and confidence loss are given a higher weight. These alterations produce accurate bounding boxes and efficient feature extraction. YOLOv3 provides detection at an excellent speed. However, it suffers from some shortcomings: expressions are affected by the external environment, and orientations/posture are not taken into account.

#### YOLOv4

YOLOv4 introduces several features that improve the learning of Convolution Neural Networks (CNNs) [29]. These include Weighted Residual Connections (WRC), Cross-Stage-Partial connections (CSP), Cross mini-Batch Normalization (CmBN), and Self-adversarial training (SAT). CSPDarknet is used as an architecture. It contains 29 convolutional layers 3 × 3, a 725 × 725 receptive field, and 27.6M parameters. Spatial Pyramid Pooling (SPP) is added on the top of this layer. YOLOv4 improves the Accuracy Precision Score and FPS of v3 by 10 to 12%. It is faster, more accurate, and can be used on a conventional GPU with 8 to 16 GB-VRAM which enables widespread adoption. New features suppress the weakness and improve on the already impressive face detection capabilities of its predecessor.

#### Faster R-CNN

Faster R-CNN is the third iteration of the R-CNN architecture. Rich feature hierarchies for accurate detection of objects and features, and semantic segmentation CNN (R-CNN) started in 2014, introducing a method of Selective Search to detect regions of interest in an image and a CNN to classify and adjust them [30]. However, it struggled with producing real-time results. The next step in its evolution was Fast R-CNN, published in early 2015: a faster model with shared computation capabilities owing to the Region of Interest Pooling technique. Finally came Faster R-CNN, the first fully differentiable model. The architecture consists of a pre-trained CNN (ImageNet) up until an intermediate layer, which gives a convolutional map. This is used as a feature extractor and provided as input to Region Proposal Network, which tries to find bounding boxes in the image. Region of Interest (RoI) Pooling then extracts features that correspond to the relevant objects into a new tensor. Finally, the R-CNN module classifies the contents in the bounding box and adjusts its coordinates to better fit the detected object. Maximum pooling is used to reduce the dimensions of extracted features. A Softmax layer and a regression layer were used to classify facial expressions. This results in Faster R-CNN achieving higher precision and lower miss-rate. However, it is prone to overfitting: the model can stop generalizing at any point and start learning noise.

## 3. Results

### 3.1. Model Parameters

YOLOv3 and YOLOv4 were given image inputs in batches of 64. Learning rate, Momentum, and Step Size were set to 0.001, 0.9, and 20000 steps, respectively. Training took 10+ hours for the former and 8+ for the latter. Faster R-CNN accepted input in batches of 32. Learning rate, Gamma, Momentum, and Step Size were set to 0.002, 0.1, 0.9, and 15000, respectively. It is the most time-consuming to train of the 3, taking 14+ hours. The confusion matrix of DarkNet-53, CSPDarkNet-53, VGG-16 trained and tested on the farm animals’ images and videos dataset using YoloV3, YoloV4 and Faster RCNN respsectively are shown in Tables S1, S2 and S3.

### 3.2. Computation Resources

YOLOv3 with its Darknet53 architecture takes the most inference time (0.0331s) compared to YOLOv4(0.27s) and Faster R-CNN (0.3s), both of which have CSPDarknet53 and VGG-16 architectures, respectively. YOLOv4 is the computationally efficient model, using 3479 MBs compared to 4759 MBs usage by YOLOv3 and 5877 MBs by Faster R-CNN. YOLOv4 trumps its two competitors when it comes to resources and efficiency, with optimal memory usage and good-enough inference time. Figure 2 shows the images of farm animals detected by WUR Wolf facial coding platform from the dataset using Faster RCNN technique.

**Figure 2.**
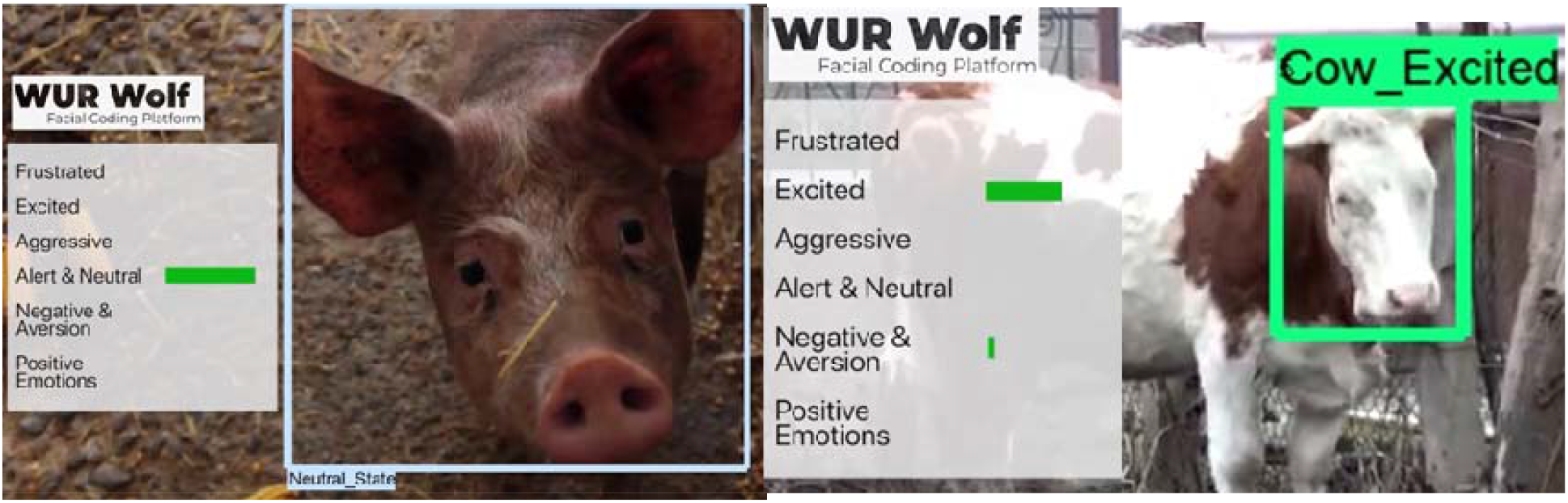
Example emotion detection results from a neutral emotional state pig, and an excited emotional state cow as determined by the WUR Wolf Facial Coding Platform.

YOLOv3 takes the least amount of time in learning most of the features than the other 2 models and the accuracy curve (Figure 3) flattens earlier as a result. Its fluctuating loss curve is a result of more repetitive predictions and slower convergence as compared to YOLOv4 and Faster R-CNN.

**Figure 3.**
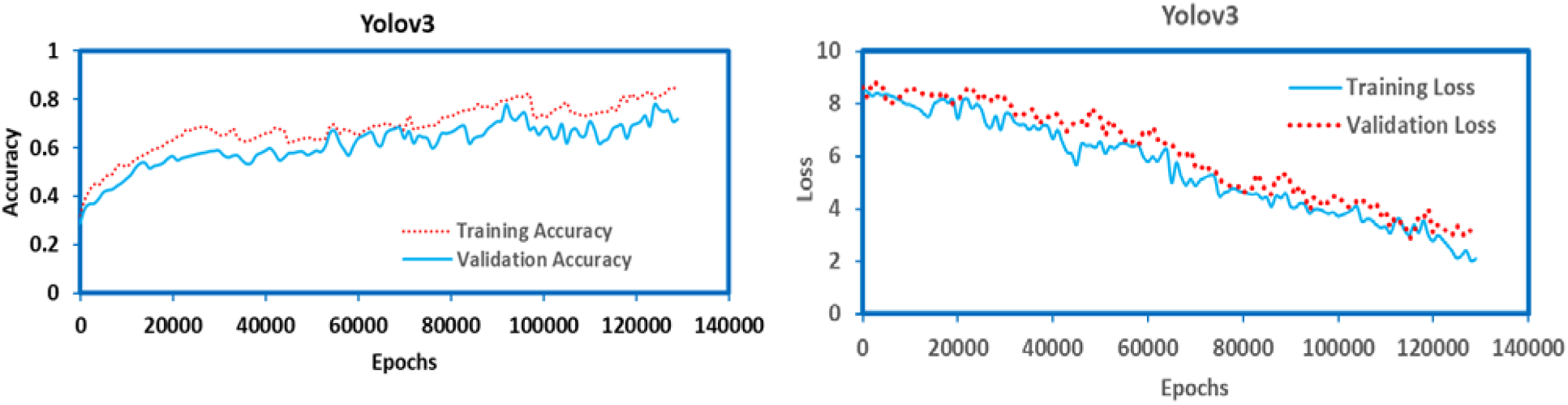
Accuracy and Error Loss for YoloV3

YOLOv4 is slower in learning than Yolov3 but achieves a higher accuracy score and a smoother loss function. Validation accuracy is very close to train accuracy as well, indicating that the model is generalizing well on unseen data (Figure 4) and would perform better in real-time than v3.

**Figure 4.**
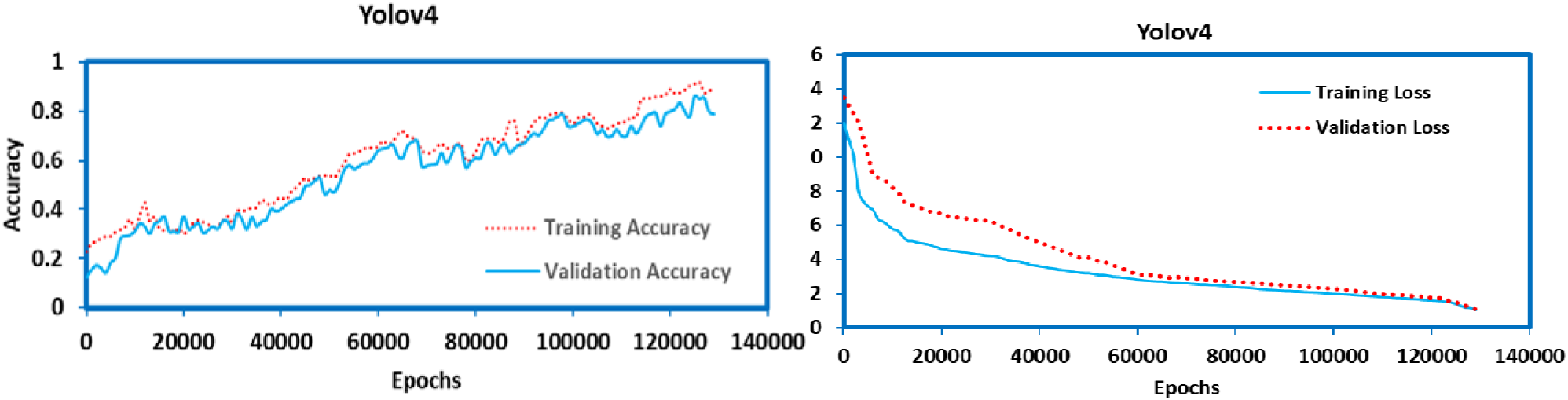
Accuracy and Error Loss for YoloV4

Faster R-CNN achieves a higher accuracy (Figure 5) score than both of the YOLO variants as well as converging quickly. However, it performs poorly in generalizing the learning as the difference between validation and train accuracy is very large at multiple times. Faster R-CNN’s accuracy score (93.11% on training and 89.19% on validation set) outperforms both YOLOv4 (89.96% on training and 86.45% on validation set) and YOLOv3 (85.21% on training and 82.33% on validation set) on these metrics. Its loss curve is also faster to converge, followed closely by v4, and v3 is the worst performer on this metric.

**Figure 5.**
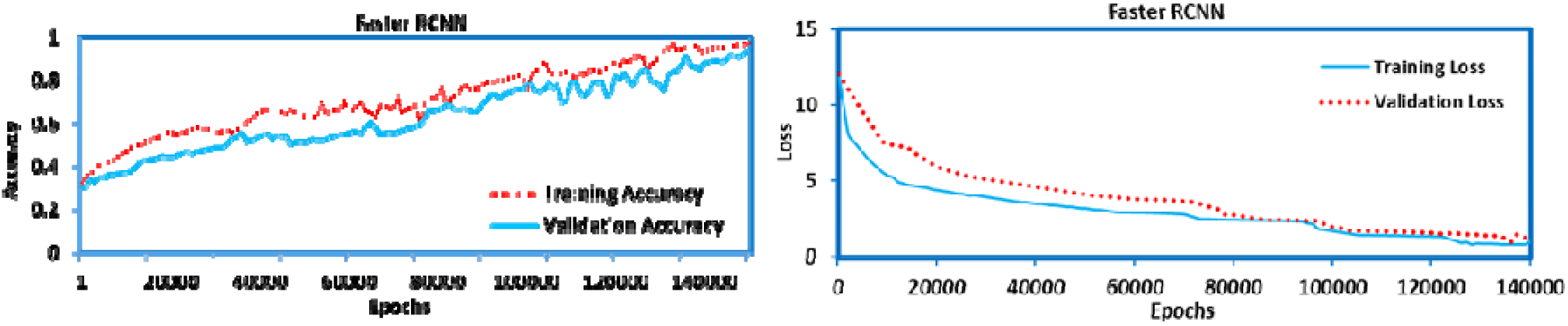
Accuracy and Error Loss for Faster RCNN

### Mean Average Precision (mAP)

The mAP score compares the actual bounding box to the detected box and returns a score. The higher the score, the more accurate is the model’s object boundary detection. YOLOv4 has a mAP score of 81.6% at 15 FPS, performing better than both the other models. YOLOv3 also performs well on this metric with a mAP score of 77.60% at 11 FPS. Faster R-CNN also provides a moderate mAP score of 75.22%; however, its processing speed is very slow at just 5 FPS. Among the 3, YOLOv4 provides the best bounding boxes at a higher speed.

### F1 Score

The annotated labels for both cows and pigs can be grouped on the basis of mental states such as positive, negative, and neutral. Analyzing model performance on these groups is useful in measuring how the model works in different contexts. F1 score is a good measure for this analysis. A Confusion Matrix tabulates the performance of a model on the dataset for which true values are known. Model results are compared against pre-set annotations, and an analysis of them reveals the performance of each model in detecting the emotion portrayed in the picture. Confusion Matrices of all three models are given in the Supplementary Reading Section alongside respective F1 scores. Negative context requires additional effort and reactions, and as a result, there are more pixels with useful information in classification. All 3 models perform return higher True Positives for such cases (Table S4, S5, S6). The average F1 scores of each of the models are as follows: 85.44% for YOLOv3, 88.33% for YOLOv4, and 86.66% for Faster R-CNN. YOLOv4 outperforms the other two in predicting emotion states for each image.

## 4. Discussion

Non-invasive technology that can assess good and poor welfare of farm animals, including positive and negative emotional states, is possible soon using the proposed WUR Wolf Facial Coding Platform. The ability to track and analyze how animals feel will be a breakthrough in establishing animal welfare auditing tools.

In this project, we evaluated the applicability of 3 deep learning-based models for determining the emotions of farm animals, Faster R-CNN, and two variants of YOLO: YOLOv3 and YOLOv4. For training the YOLO3 and YOLO4 algorithms, we used the darknet framework. YOLO4 has the CSPDarknet53, while YOLOv3 has the Darknet53. Because of the differences between the backbones, Yolov4 is faster and provides more accurate results for real-time applications.

Demonstration and results of emotion detection of cows and pigs using Faster RCNN (Figure 2) is shown in the attached supplementary video [V1]. Faster RCNN is suitable for mobile terminals where there is a lack of hardware resources in facial expressions recognition [31]. If speed (time for data processing) is the deciding factor, then YoloV4 is a better choice than Faster RCNN. Due to the advantage of the network design, large variations in the dataset composed of facial images and videos with complex and multiscale objects is better analyzed by the two-stage Faster RCNN method. Hence, for higher accuracy in the results of emotion detection, Faster RCNN is recommended over YoloV4. In on-farm conditions where there may be a lack of equipment related to strong data processing ability, Faster RCNN would be a good choice. Technological advances in the field of animal behavior are a huge step in improving humans’ understanding of the animals they share this world with, but there is still room to grow.

No facial recognition software created for animals is 100% accurate yet, and so far, only a few common species and non-human primates have had this software modified to identify their physical features. Animal species that are not identified as mammals are minimally expr
essive and have not been tested with facial recognition software for the study of their emotions. One study even brought up the consideration that animals may be able to suppress emotional expression, much like people do in situations where it is socially appropriate to express only certain emotions [25]. There are many questions related to animal emotional expression that have yet to be answered, but there is a good chance that the advancement and implementation of facial recognition software will lead scientists to those answers in the future.

## 5. Conclusions

The detailed analysis of the performance of the 3-machine learning python-based models shows the utility of each model in specific farm conditions and how they compare against each other. YOLOv3 learns quickly but gives random predictions and fluctuating losses. Its next iteration, YOLOv4, has improved considerably in multiple regards. If the aim is to balance higher accuracy with faster response and less training time, YOLOv4 works best. If the speed of training and memory usage isn’t a concern, the 2-staged Faster R-CNN method performs well and has a robust design for predicting different contexts. The output is accurate, and overfitting is avoided. There is no one-size-fits-all model, but with careful consideration, the most efficient and cost-effective methods can be selected and implemented in automating the facial coding platform for determining farm animal emotions. Facial features and expressions of farm animals provides only one-dimensional aspect of the affective states. Due to the advent of Artificial Intelligence and sensor technologies, in the near future multi-dimensional models of mental and emotional affective states will emerge in the form of measuring behavioural patterns, combined track changes in farm animal postures and behavioural changes with large-scale neural recordings.

## Supplementary Materials

The following are available online at www.mdpi.com/xxx/s1, Figure S1: title, Table S1: Statistical Analysis Results of the Images, Video S1: Demonstration of the WUR Wolf Facial Coding Platform.

## Author Contributions

Conceptualization, S.N.; methodology, S.N.; algorithm development & coding, S.N.; validation, S.N.; formal analysis, S.N.; investigation, S.N.; writing—original draft preparation, S.N.; writing—review and editing, S.N.; supervision, S.N.; project administration, S.N; All authors have read and agreed to the published version of the manuscript.

